# Performances of NeuMoDx, a random-access system for HBV-DNA and HCV-RNA quantification

**DOI:** 10.1101/2020.10.14.340398

**Authors:** Juliette Besombes, Charlotte Pronier, Charles Lefevre, Gisèle Lagathu, Anne Maillard, Claire Grolhier, Vincent Thibault

## Abstract

Viral loads (VL) monitoring for hepatitis B and C is essential to evaluate disease progression and treatment response. Automated, random-access rapid systems are becoming standard to provide reliable VL to clinicians. The aim of this study was to evaluate the analytical performances of the recently launched NeuMoDx^™^ for HBV-DNA and HCV-RNA quantification. Clinical samples routinely quantified on the Beckman-Veris system were either retrospectively (frozen samples; HBV n=178, HCV n=249), or in parallel (fresh primary tubes; HBV n=103, HCV n=124) tested using NeuMoDx^™^. Linearity range was assessed on serial dilutions of high tittered plasmas containing different genotypes for HBV (A-E, n=10) and HCV (1a-b, 2-5, n=12). Overall test failure, mostly internal control amplification failure, was 2.3% and was not influenced by matrix types. For HBV-VL, Kappa agreement was 74%, with 27 (12.6%) discrepancies. Correlation between HBV assays on 72 quantified samples by both methods was excellent (r=0.963) with a mean bias (NeuMoDx^™^-Veris) of 0.21 log IU/mL. For HCV-VL, Kappa agreement reached 94%, with 9 (2.8%) discrepancies. The r-correlation factor between assays on 104 samples was 0.960 with a mean bias of −0.14 log IU/mL (NeuMoDx^™^-Veris). Serial dilutions confirmed the claimed linear ranges for all HBV and HCV genotypes. The mean turnaround time was 72’ [55-101] for HBV and 96’ [78-133] for HCV. These results obtained on the NeuMoDx^™^ confirmed the overall good functionality of the system with a short turn-around-time, full traceability and easy handling. These results on HBV- and HCV-VL look promising and should be challenged with further comparisons.

## Background

Hepatitis B virus (HBV) and Hepatitis C virus (HCV) are the most common chronic viral infections and remain an important global health problem. It is estimated that more than 250 million people are living with chronic HBV infection while approximately 70 million people are chronically infected with HCV (1). These chronic infections may lead to chronic liver diseases, such as cirrhosis, and to primary liver cancer (hepatocellular carcinoma). The incidence and prevalence of these infections have notably decreased due to universal birth HBV vaccination programs (2, 3) and to new Direct Acting Anti-viral (DAA) drugs used for HCV infection (4). But HBV and HCV are still a main cause of deaths worldwide, especially in low to middle income countries, and are responsible for 96% of the mortality from viral hepatitis (1).

Serological markers alone are not sufficient to document an infection. Indeed, anti-HCV antibody detection does not differentiate an active from a past infection, the former being necessarily confirmed by viral RNA detection (5). Determination of HBV and HCV viral load (VL) in serum or plasma is essential to diagnose an acute or chronic infection, to establish the prognosis, to decide the course of treatment and to monitor the virologic response to antiviral therapy (6, 7). The ultimate objective for the patients is to achieve an undetectable VL in both cases. The risk of disease progression is reduced when a sustained reduction of HBV-DNA to undetectable levels is achieved whatever the course of the infection (6). In a similar way, remission from an HCV infection following treatment by DAA is defined by the achievement of a sustained virological response (SVR) that is an undetectable HCV RNA level 12 weeks (SVR12) or 24 weeks (SVR24) after treatment completion (7). Thus, international clinical practice guidelines recommend the use of sensitive nucleic acid amplification technologies for HBV-DNA or HCVRNA detection (6–9). Assays with lower limit of detection ≤10 International Units (IU)/mL for HBV and ≤15 IU/mL for HCV RNA are strongly recommended (6, 7).

Fully automated, random-access, closed systems that integrate the entire molecular diagnostic process from “sample to result” are routinely used for this purpose (10, 11). They include reagent storage, specimen preparation, nucleic acid extraction, real time polymerase chain reaction (PCR) setup, amplification, and detection, as well as result analysis and reporting, on a single platform. Among many advantages, they usually offer high throughput, full traceability, minimal cross-contamination risk and high safety standard for end users.

The NeuMoDx^™^ 96 Molecular System is a recently launched fully automated, continuous random-access analyser, which integrates magnetic particle affinity capture and real time PCR chemistry in a multi-sample microfluidic cartridge. The aim of this study was to evaluate the analytical performances of the NeuMoDx^™^ for HBV-DNA and HCV-RNA quantification. We used a panel of specimens, including different HBV and HCV genotypes, with a wide range of VL values as determined on a routine basis with the VERIS MDx System (Beckman-Coulter) to compare the concentrations obtained with the new system.

## Materials and Methods

### HBV and HCV viral load assays (table 1)

The NeuMoDx^™^ Molecular System is a fully automated sample to result molecular diagnostic system: it integrates the extraction, purification, quantification, and result interpretation of infectious disease nucleic acid targets. It is based on quantitative real time PCR performed in patented universal microfluidic cartridges. The plasma input volume is 1.0 mL. An exogenous Sample Processing Control is incorporated in the Extraction Plate. The NeuMoDx^™^ HBV Quant Assay targets the highly conserved sequences in the region encoding X protein and pre Core protein in the HBV genome. Across genotypes, the dynamic range of quantification for the HBV assay is 7.6 to 1.05E+09 IU/mL (0.88 to 9.02 log IU/mL). The NeuMoDx^™^ HCV Quant Assay targets the highly 5’UTR conserved sequences in the HCV genome. Across genotypes, the dynamic range of quantification for the HCV assay is 7.7 to 1.6E+08 IU/mL (0.9 to 8.2 log IU/mL). The system is CE and FDA approved.

**Table 1:** Summary of HBV and HCV assays characteristics.

| Systems | Reagents | Target Region | Limit of quantification (IU/mL) | Input volume<br>(+dead volume) | Acquisition time |
| --- | --- | --- | --- | --- | --- |
| NeuMoDx™ 96 | HBV assay | Encoding HBx and Pre C proteins | 7.6 |  | 60' |
|  |  |  |  | 1 mL |  |
|  | HCV assay | Highly conserved | 7.7 |  | 80' |
| Veris MDx<br>(Beckman Coulter) | HBV assay | S gene | 10 |  | 70' |
|  |  |  |  | 1 mL |  |
|  | HCV assay | 5'UTR | 12 |  | 105' |

The VERIS MDx System (Beckman Coulter) is also a random access integrated automated nucleic acid extraction and real-time PCR system that uses a plasma input volume of 1.0 mL. Quantification is based on the amplification of S gene for HBV and 5’ non-coding region for HCV. The dynamic range of quantification of the HBV assay is 10 IU/mL to 1,000,000,000 IU/mL (1–9 log IU/mL) and 12 to 100,000,000 IU/mL (1–8 log IU/mL) for the HCV assay. An internal control is included in each test to monitor process variations. The acquisition time is 70 min for 1 patient after loading a sample tested for HBV and 105 min for HCV. The system is CE-marked.

These tests establish traceability to the 4^th^ WHO International Standard for HBV assay and to the 5^th^ for HCV assay.

External quality controls (positive and negative) were passed daily prior to start of testing for both assays on both systems.

### Sample collection

The study included 281 samples for HBV assay and 373 samples for HCV assay from patients of the University Hospital Pontchaillou of Rennes, France. Human whole blood was collected in sterile blood collection tubes containing Ethylenediaminetetraacetic acid (EDTA) as anticoagulant for the preparation of plasma, which was stored at −80°C after centrifugation. They were collected during routine VL measurements and initially quantified on the Veris MDx system. Then, they were either retrospectively (frozen samples; n=178 HBV, n=249 HCV) or in parallel (fresh primary tubes; n HBV=103, n HCV=124) tested using NeuMoDx^™^ specific reagents (Fig.1).

**Figure 1:**
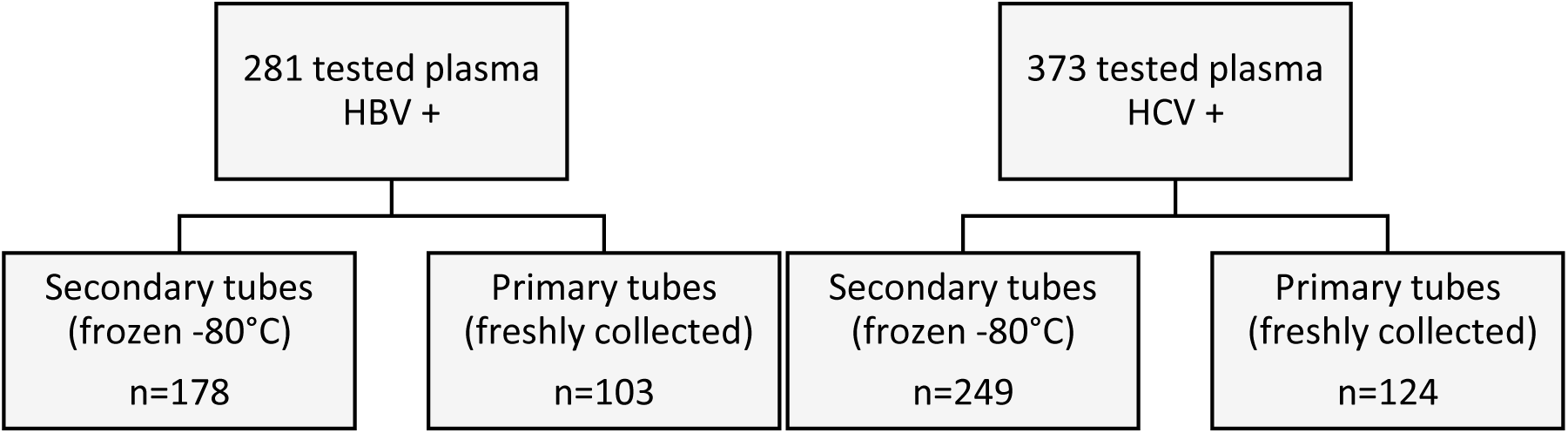
Flowchart of sample processing.

### Linearity

The linearity of quantification by the NeuMoDx^™^ was assessed on serial dilutions of high tittered plasmas (around 7 log IU/mL) containing different genotypes, for both HBV and HCV assays in order to cover the assay full quantification ranges. Nine Ten HBV positive plasmas (genotype A to E) and 12 HCV positive plasmas (genotype 1a, 1b, 2, 3, 4, 5) were selected among frozen samples from the plasma collection. Tenfold dilutions were performed in HBV and HCV negative human plasmas supplied by the Blood Transfusion Center. The measured values were compared to the expected values calculated using initial Veris VL.

### Reproducibility

To assess inter-assay reproducibility, a set of two HBV and HCV daily controls, including a low-positive and a high-positive controls, were tested eleven times (on different days).

### Statistical analysis

All VL values were transformed to log10 format for further statistical analysis. Passing-Bablok regression was performed to evaluate the correlation between the NeuMoDx^™^ and the Veris while Bland–Altman analysis was used to analyze the concordance and mean difference between assays. Overall agreement was measured by weighted Cohen’s kappa coefficient. Means of all differences and standard deviations (SDs) were calculated. The 95% limits of agreement between assays were determined as the mean ± 1.96 SD. Statistical significance was set at *P* < 0.05. (Analyse-it v5.30 https://analyse-it.com/).

## Results

### Linearity

### HBV assay

Statistical analysis was performed on the 47 dilutions out of 67 (Fig.2), with VL ranging from 1.33 log IU/mL to 7.51 log IU/mL covering the linearity range of the NeuMoDx^™^ HBV assay. Correlation between HBV-DNA levels measured by the NeuModx^™^ HBV assay and the expected values was excellent (r=0.99, IC 95%: 0.91 to 1.1, linear regression equation: y = 0,690 + 0,985 x) (Fig.2A). A modest bias (mean bias = 0.61 log IU/mL, IC 95%: −0.14 to 1.4) was observed between NeuMoDx^™^ HBV and the expected values (Fig.2B). These differences seemed to be more important in lower values, close to the quantification threshold of the technique (<1 log IU/mL). When comparing with the expected values, a clinically significant difference in quantification (>0.5 log) was observed throughout the linearity range of the assay. No influence of genotype on VL quantification was identified.

**Figure 2:**
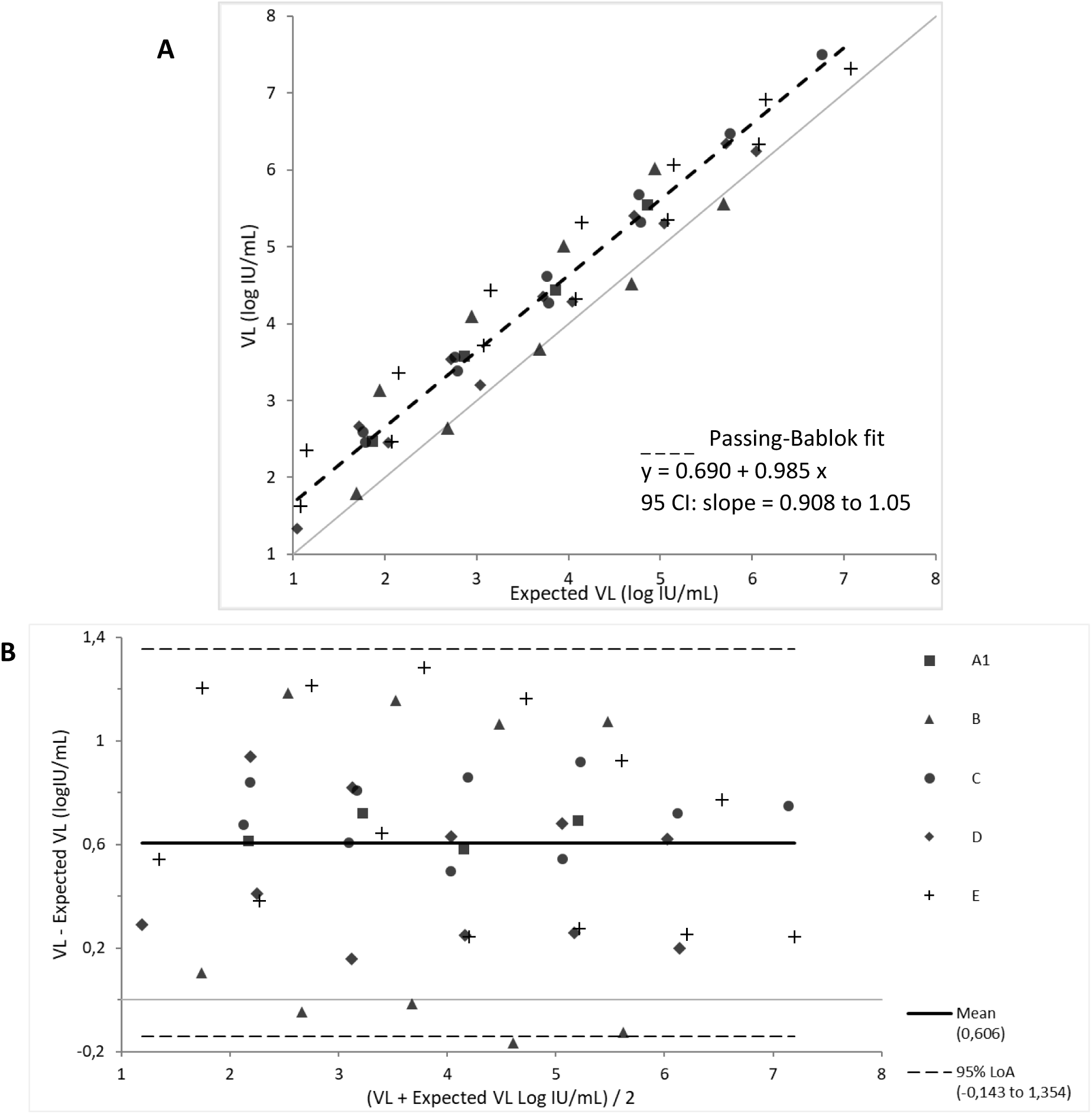
Passing Bablok regression and Bland-Altman plot analysis of HBV-DNA levels as determined by the NeuMoDx HBV assay on 47 samples obtained after dilution of 9 clinical specimens from patients infected with HBV genotypes A (n =1), B (n =2), C (n = 2), D (n = 2) and E (n = 2). (A) Passing Bablok regression of NeuMoDx HBV assay versus expected values. Dashed line represents the regression line. (B) Bland-Altman plots of NeuMoDx HBV versus expected values. On the Bland-Altman graph, the bold and dashed lines represent the mean and the 95% limits of agreement, respectively.

#### HCV assay

Statistical analysis was performed on the 52 dilutions out of 84 (Fig.3), with a VL ranging from 1.1 log IU/mL to 6.19 log IU/mL covering the linearity range of the NeuMoDx^™^ HCV assay. Correlation between HCV RNA levels measured by the NeuModx^™^ HCV assay and the expected HCV RNA levels with the Veris was very good (r=0.89, IC 95%: 0.81 to 0.97, linear regression equation: y = 0,196 + 0,893 x) (Fig.3A). A little bias (mean bias = −0.095 log IU/mL, IC 95%: −1.11 to 0.92) was observed between NeuMoDx^™^ HCV and the expected viral loads (Fig.3B). This dispersion was more important in lower values, close to the quantification threshold of the technique (<1 log IU/mL). No influence of genotype on VL quantification was identified.

**Figure 3:**
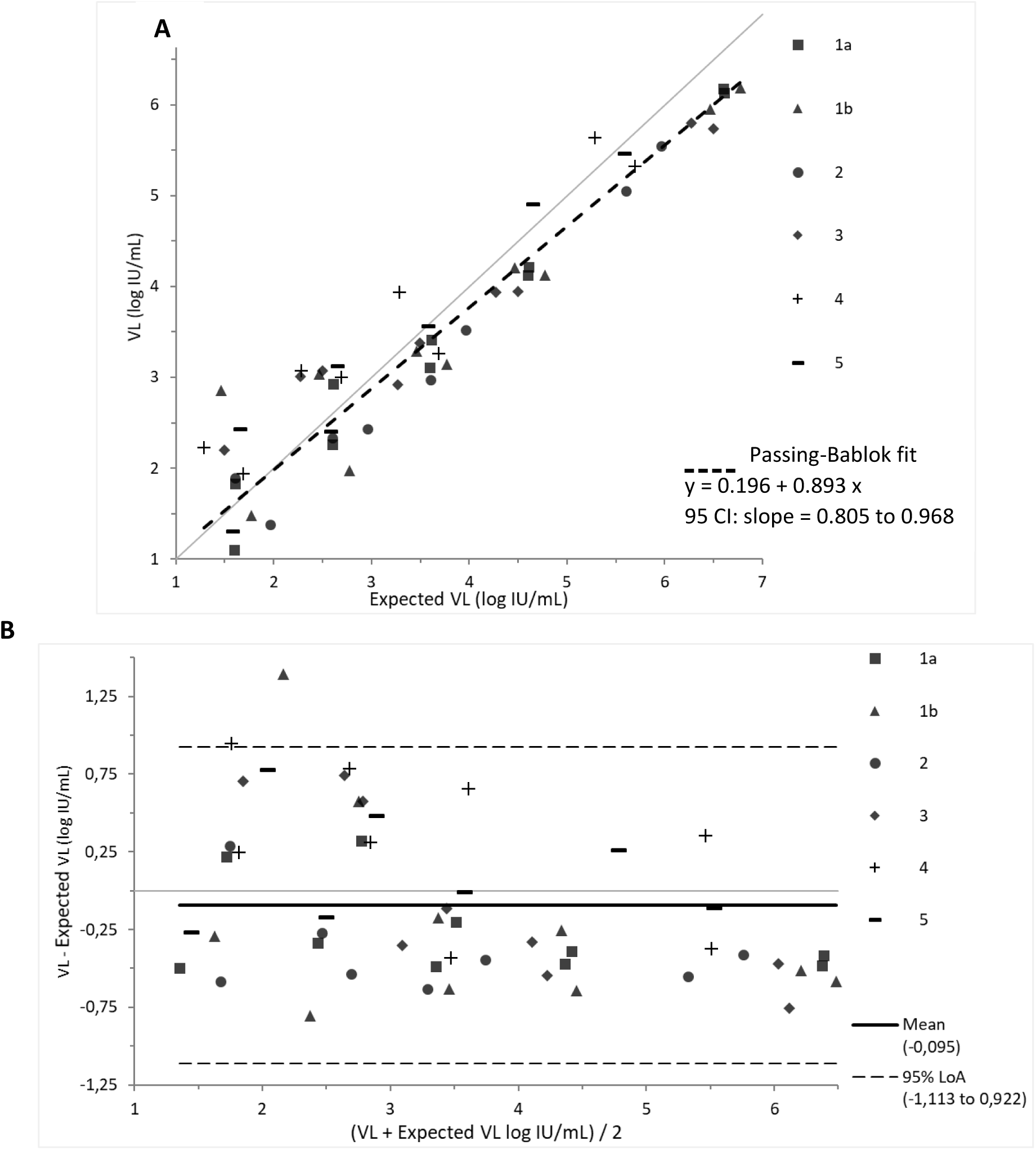
Passing Bablok regression and Bland-Altman plot analysis of HCV RNA levels as determined by the NeuMoDx HCV assay on 52 samples obtained after dilution of 12 clinical specimens from patients infected with HCV genotypes 1a (n =2), 1b (n =2), 2 (n = 2), 3 (n = 2), 4 (n = 2) and 5 (n=2). (A) Passing Bablok regression of NeuMoDx HCV assay versus expected values. Dashed line represents the regression line. (B) Bland-Altman plots of NeuMoDx HCV versus expected values. On the Bland-Altman graph, the bold and dashed lines represent the mean and the 95% limits of agreement, respectively.

#### Reproducibility

For each assay, a low and a high control specimens were measured eleven times by different users and over several days. The inter-assay coefficients of variation (CV) for high and low HBV-DNA levels were 1.48% and 4.86% respectively. For high and low HCV-RNA levels, they were 1.77% and 6.81% respectively.

### Performances

#### HBV assay

HBV-DNA levels were measured in parallel with both NeuMoDx^™^ and Veris systems in 281 samples from patients infected with HBV genotypes A to E. Overall test failure, corresponding mostly to internal control amplification failure was 2.3% and was not influenced by the matrix types (fresh or frozen sample). The HBV statuses obtained with the different tests are described in Table 2. Among the 102 positive samples with the Veris assay, 70.6% were also positive with the NeumoDx^™^ assay. None of the negative samples with the Veris assay was detected positive with the NeumoDx^™^ assay but 70.6% of detected samples on the Veris were negative on the NeumoDx^™^. Seventy-two samples were quantified with the NeuMoDx^™^ and the Veris assays with a quantification range of 0.74 (8 IU/mL) – 8.82 log IU/mL and of 1.08–8.75 log IU/mL respectively. The overall Kappa qualitative agreement was 74%, with 27 (12.6%, 16 “detected” and 11 quantified) discrepancies (Veris Positive while NeuMoDx^™^ negative); discrepant sample VL ranged from 1.1 to 2.6 log IU/mL. Correlation between both HBV assays on 72 quantified samples was excellent (r=0.967, linear regression equation: y = 0.052 + 1.03x) (Fig.4A) with a mean bias (NeuMoDx^™^-Veris) of 0.21 log IU/mL (95% CI: 0.08 to 0.33) (Fig.3B).

**Figure 4:**
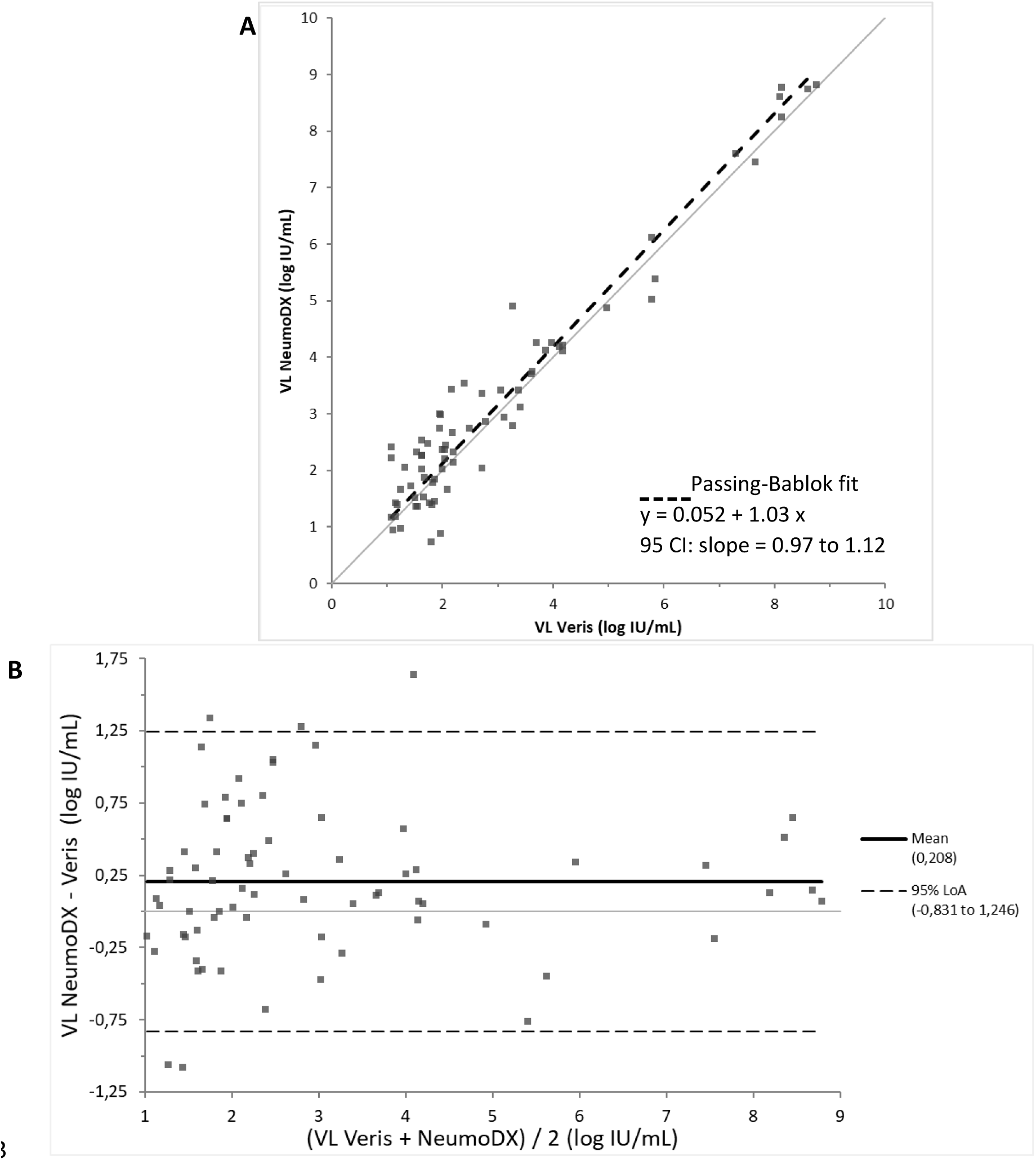
Passing Bablok regression and Bland-Altman plot analysis of HBV-DNA levels measured by the NeuMoDx and Veris HBV assays in 72 samples from patients infected with HBV. (A) Passing Bablok regression of NeuMoDx HBV assay versus expected values. Dashed line represents the regression line. (B) Bland-Altman plots of NeuMoDx HBV versus expected values. On the Bland-Altman graph, the bold and dashed lines represent the mean and the 95% limits of agreement, respectively.

**Table 2:**
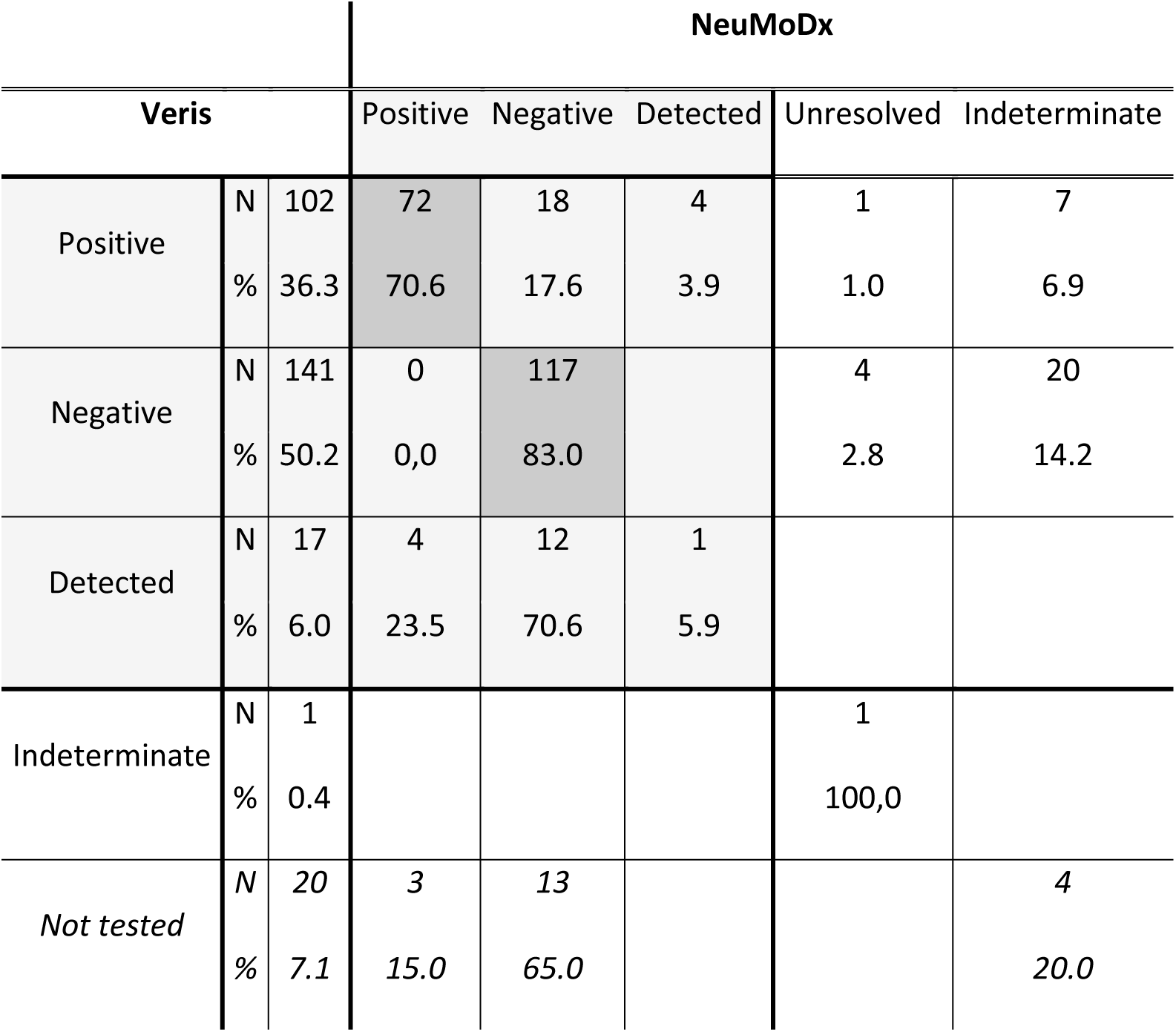
Comparison between NeumoDx^™^ and Veris (reference) HBV assays among the 281 selected samples. Possible statuses are: positive, negative, detected, unresolved (usually defines an amplification failure) or indeterminate (usually indicates a sample processing failure like low volume).

|  |  |  | NeuMoDx |  |  |  |  |
| --- | --- | --- | --- | --- | --- | --- | --- |
| Veris |  |  | Positive | Negative | Detected | Unresolved | Indeterminate |
| Positive | N | 102 | 72 | 18 | 4 | 1 | 7 |
|  | % | 36.3 | 70.6 | 17.6 | 3.9 | 1.0 | 6.9 |
| Negative | N | 141 | 0 | 117 |  | 4 | 20 |
|  | % | 50.2 | 0,0 | 83.0 |  | 2.8 | 14.2 |
| Detected | N | 17 | 4 | 12 | 1 |  |  |
|  | % | 6.0 | 23.5 | 70.6 | 5.9 |  |  |
| Indeterminate | N | 1 |  |  |  | 1 |  |
|  | % | 0.4 |  |  |  | 100,0 |  |
| Not tested | N | 20 | 3 | 13 |  |  | 4 |
|  | % | 7.1 | 15.0 | 65.0 |  |  | 20.0 |

#### HCV assay

HCV-RNA levels were measured in parallel with both NeuMoDx^™^ and Veris systems in 373 samples from patients infected with HCV genotypes 1a, 1b, 2 to 5. Overall test failure, corresponding mostly to internal control amplification failure was 3.2% and was not influenced by the matrix type (fresh or frozen sample). The HCV statuses obtained with the different tests are described in Table 3. None of the positive sample with the Veris assay was negative with the NeumoDx^™^ assay. One hundred and four (94.5%) samples were quantified with the NeuMoDx^™^ and the Veris assays with a quantification range of 1.32–7.44 log IU/mL and of 1.11–7.61 log IU/mL, respectively. Among the 223 negative samples with Veris assay, 87.4% were also negative with the NeumoDx^™^ assay. The overall Kappa qualitative agreement reached 94%, with 9 (2.8%, 6 quantified) discrepancies (Veris Negative while NeuMoDx^™^ Positive). Among those, 8 HCV-VL were detected positive (HCV-VL from 17 to 71 IU/mL) on NeuMoDx^™^ while negative on Veris. The r-correlation factor between both HCV assays on 106 samples was also excellent (r=0.960, linear regression equation y= 0.16 + 0.94x) (Fig.4A) with a mean bias (NeuMoDx^™^-Veris) of −0.14 log IU/mL (95% CI: −1.15 to 0.86) (Fig.4B).

**Table 3:**
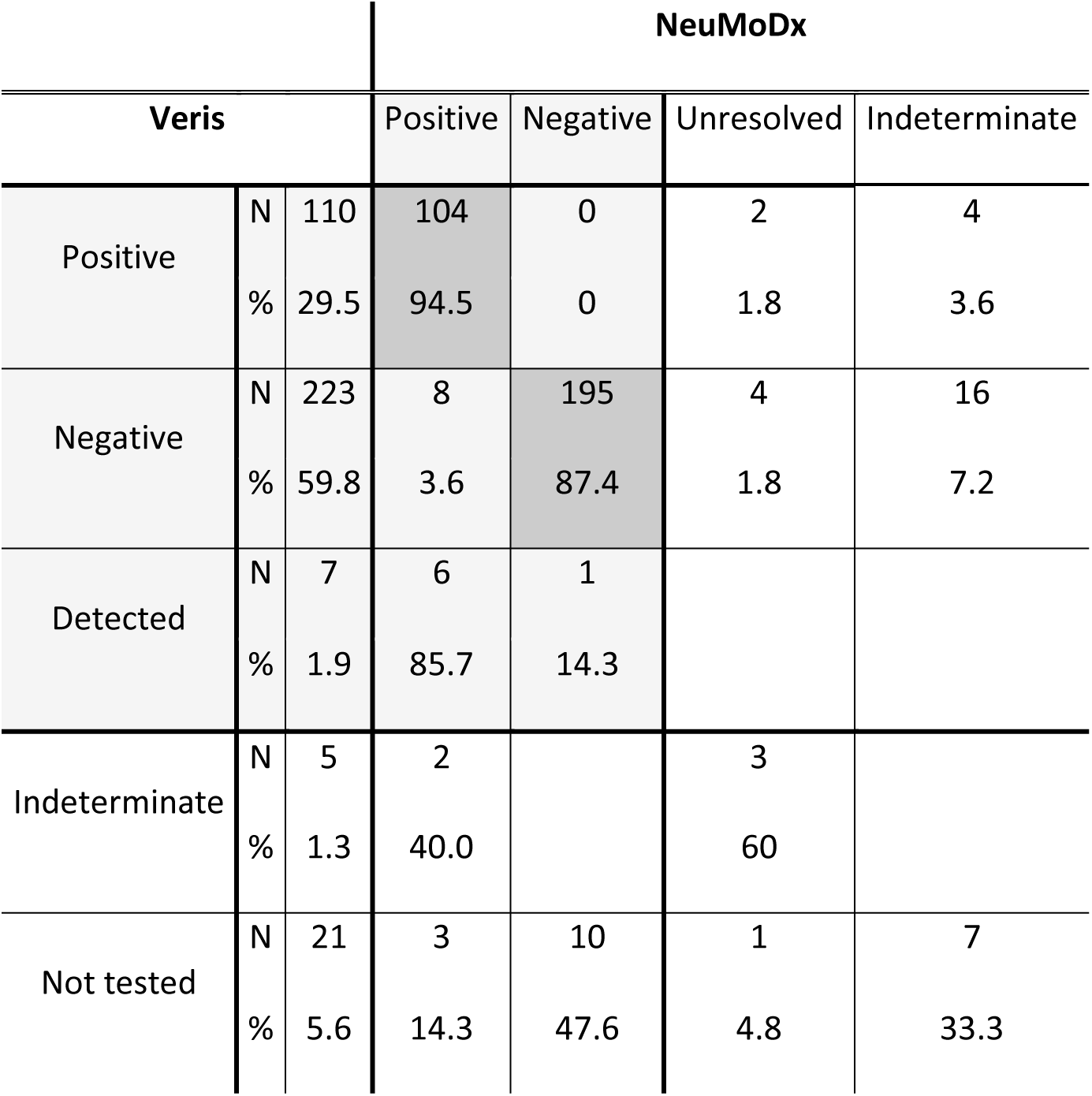
Comparison between NeumoDx^™^ and Veris (reference) HCV assays among the selected samples. Possible statuses are: positive, negative, detected, unresolved (usually defines an amplification failure) or indeterminate (usually indicates a sample processing failure like low volume).

**Figure 5:**
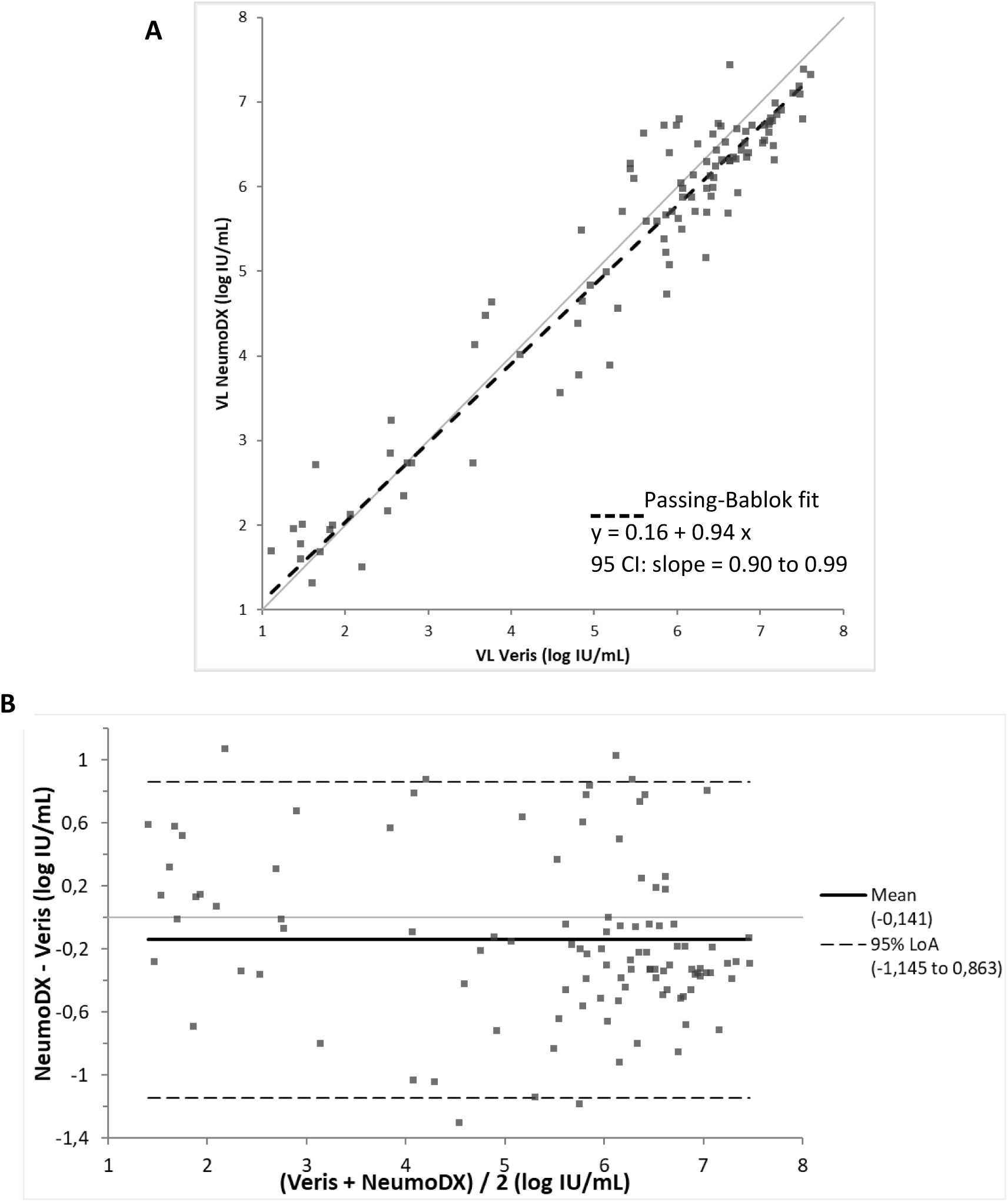
Passing Bablok regression and Bland-Altman plot analysis of HCV RNA levels measured by the NeuMoDx and Veris HCV assays in 104 samples from patients infected with HCV. (A) Passing Bablok regression of NeuMoDx HCV assay versus expected values. The dashed line represents the regression line. (B) Bland-Altman plots of NeuMoDx HCV versus expected values. On the Bland-Altman graph, the bold and dashed lines represent the mean and the 95% limits of agreement, respectively.

#### Turnaround time

On NeuMoDx^™^ the mean turnaround time was 72’ [55-101] for HBV and 96’ [78-133] for HCV and depended on the number of concomitantly loaded samples on the system.

## Discussion

As recommended by international guidelines, real time PCR has become the gold-standard to provide reliable quantification of HBV and HCV-VL to clinicians in reasonable time. This measurement is necessary to diagnose chronic HBV or HCV infection, to make decision for therapeutic intervention and to monitor antiviral treatment responses and resistance (6, 7). Fully automated and continuous random-access analysers are routinely used and offer a fast turn-around time that also allows a rapid screening of an organ, tissue or cell donor prior to transplantation. Such systems are usually simple to use and accessible to individuals with no background in molecular biology or virology.

The NeuMoDx^™^ is a new launched fully automated continuous random-access molecular diagnostic platform with full traceability. The specificities of this system are its process based on microfluidic and its use of dried reagents that are stable at room temperature. In this study, we evaluated the performance of the system as a routine system.

The Veris MDx System is suitable for high-throughput HBV-DNA and HCV-RNA monitoring in large hospital laboratories and was well suited as a routine system for our lab (10). Both HBV and HCV NeuMoDx assays were compared to those performed on the Veris. We found that both NeuMoDx assays have a good linearity over the claimed quantitation ranges. Assays reproducibility was good, particularly for high viral-load controls; the CV of both HBV and HCV assays for controls at 5 log IU/mL was below 2%. At lower viral loads, the HBV assay CV (4.86%) appeared better than the one observed on the Veris or Genexpert assays but worse than what reported on the Cobas AmpliPrep/Cobas TaqMan HBV test (10–12). We noticed a more pronounced dispersion in the low VL values for the HCV test that may require further investigation even though it remained acceptable. This CV (6.81%) for low HCV-RNA levels is similar to those observed with the Genexpert or Veris assays but better than the one with VERSANT 1.0 assay (13–15). Higher CV at the lower end of quantification is commonly seen with other systems and we cannot affirm from our study whether this observation was more important on NeuMoDx.

The concordance with the Veris was better for the HCV (k=0.94) than for the NeuMoDx HBV assay (k=0.74). Yet, an overall very good quantification correlation was observed between NeuMoDx and Veris for both HBV (r=0.929) and HCV (r=0.960) assays. Even though the NeuMoDx assays possess performances comparable to those provided by the Veris, discrepant quantifications were mostly observed on low VL samples. Indeed, 8 HCV-VL were detected positive on NeuMoDx^™^ while negative on Veris and conversely 27 HBV-VL were detected positive on Veris while negative on NeuMoDx^™^. The mean bias estimated for both HBV (m=0.21 log IU/mL) and HCV (m=-0.14 log IU/mL) assays should not translate into any major clinical impact. Indeed, for patient’s follow-up, a variation below 0.5 log IU/mL is usually not considered as clinically relevant (16). Yet, these subtle differences confirmed the importance to follow an infected patient with the same technique over-time, in order not to miss a relapse or a potential recontamination with HCV.

As previously frozen samples but also freshly drawn sample were run, no interference from freezing and thawing were noticed. We did not observe any difference of quantification according to genotype in our set of samples and linearity was similar whatever the genotype. Subtle observed differences could possibly be explained by differences in the extraction methods or in primer and probe sequences. Indeed, the NeuMoDx^™^ HBV assay targets regions encoding HBx and preC proteins whereas the Veris HBV assays targets the S gene. Mutation of the target region may influence the result between two techniques (17). A more exhaustive study of the different genotypes and phenotypes HBeAg positive or negative (Precore and Core mutants) should be performed to confirm their absence of influence on the quantification results.

Finally, the NeuMoDx^™^ can deliver results in roughly 1.2 h for HBV test and 1.6 h for HCV test, a fast turn-around time which is useful for emergencies such as in a transplantation context. The volume required for testing is comparable to that required in other molecular assays such as the Veris or GeneXpert (10, 14).

The automate was simple to use without any troubleshooting. The system works at room temperature due to the use of dry reagents (NeuDry^™^ technology). This enable waste reduction compared to traditional reagents and an easier storage. Moreover, the absence of cooling system reduces noise emission and energy consumption. These specificities could make this automated system suited for use in countries with limited resources, particularly in Africa where the prevalence of chronic HBV infection is higher than in Europe (6.1% vs 1.6%) (1).

These first results obtained on the NeuMoDx^™^ confirmed the overall good functionality of the system with a short turnaround time, simple training, full traceability and easy handling. Preliminary results on HBV- and HCV-VL look promising and should be challenged with further comparisons.

